# Scopolamine blocks context-dependent reinstatement of fear responses in rats

**DOI:** 10.1101/2021.02.24.432279

**Authors:** Laura M. Vercammen, Adrian C. Lo, Rudi D’Hooge, Bram Vervliet

## Abstract

Return of fear poses a problem for extinction-based therapies of clinical anxiety. Experimental research has discovered several pathways to return of fear, one of which is known as reinstatement. Here, we evaluated in rats the potential of scopolamine, a non-selective muscarinic receptor antagonist that is also safe for use in humans, to prevent the reinstatement of extinguished fear. We conducted three experiments with a total sample of 96 female rats. All rats went through a fear acquisition session (tone-shock pairings, CS-US), followed by two extinction sessions (CS only) and a post-extinction fear memory test. Twenty-four hours later, rats were placed in the same or a different context from extinction and received two unsignaled foot shock (US) presentations. On the following day, CS-evoked freezing returned when the reinstating USs had occurred in the same context compared to a different context (context-dependent reinstatement, Experiment 1). Systemic administration of scopolamine before or after the reinstating USs blocked the return of CS-evoked freezing on the following day (Experiments 2 and 3). Our findings suggest that administering scopolamine around the time of an aversive experience could prevent relapse of extinguished fears in humans.

## Introduction

Anxiety disorders are a class of prevalent and costly mental health disorders that have excessive fear as their main symptom (Kessler et al., 2005; Gustavsson et al., 2011). The treatment of choice is exposure therapy, which aims to reduce excessive fears by gradually exposing patients to their fear-evoking stimuli in the absence of the feared outcome (Craske et al., 2014). This technique is similar to Pavlovian fear extinction, which involves the repeated presentation of a conditional stimulus (CS) in the absence of the aversive unconditional stimulus (US) it was previously paired with. As in exposure therapy, these repeated presentations cause a gradual decline in the frequency and magnitude of the conditional fear reactions (CR; McNally, 2007). Extinction learning does not erase the original excitatory CS-US association, but instead leads to a new inhibitory CS-noUS association that suppresses the fear CR to the CS (Bouton, 2002). However, fear often returns after extinction, resulting in unacceptably high relapse rates after initially successful exposure therapy (Vervliet et al., 2013). Developing new strategies to prevent return of fear is thus essential for attenuating relapse in anxiety disorders.

Reinstatement is one mechanism of return of fear (Bouton, 2002). In the laboratory, reinstatement is evoked by unsignaled presentations of the aversive US, after completion of the CS extinction phase. Return of fear is measured subsequently in a CS-alone test (McNally, 2007). This effect is context-dependent, i.e., unsignaled US presentations in a different context do not elicit a return of fear for the CS when tested back in the original extinction context (Bouton & Bolles, 1979). This suggests that in normal reinstatement, the unsignaled US presentations establish a context-US association that retrieves the CS-US association when this context is also present during the CS-alone test (Bouton & Bolles, 1979). Thus, interventions that have the potential to prevent context conditioning during reinstatement could prevent return of fear.

It is well-established that the cholinergic system is involved in learning and memory processes, and in particular in the acquisition or encoding of novel information (Klinkenberg & Blokland, 2010; Hasselmo & McGaughy, 2004). Scopolamine, a non-selective muscarinic receptor antagonist, has been frequently used to assess the effects of cholinergic blockade on learning and memory, and is a potent amnestic agent (Klinkenberg & Blokland, 2010). Previous studies have repeatedly shown that scopolamine administration prior to training abolishes context conditioning in rats (Anagnostaras et al., 1995; Anagnostaras et al., 1999; Luyten et al., 2017). Hence, the current study was set up to evaluate the ability of scopolamine to prevent return of fear, by thwarting context conditioning during reinstatement. Most pre-clinical studies have focused on optimizing fear extinction learning and consolidation to prevent later relapse (Walker et al., 2002; Woods & Bouton, 2006). We took a different approach by targeting the encoding process during reinstatement instead. This could open a new psychopharmacological opportunity to prevent relapse of anxiety in patients in remission, when they are exposed to impactful aversive events.

The current study consists of three experiments. In the first experiment (EXP1), we established the standard effect of context-dependent reinstatement. In the second experiment (EXP2), we showed that scopolamine administered prior to reinstating US presentations blocks the return of CS-evoked fear. In the third experiment (EXP3), we observed the same effect when scopolamine was administered immediately after the reinstating US presentations.

## Methods and Materials

### Subjects

Eight week-old female Sprague-Dawley rats (total number of rats over the three experiments, *N* = 96) were purchased from Elevage Janvier (Le-Genest-Saint-Isle, France). The rats were group-housed in standard animal cages (*N* = 6). All animals were kept under conventional laboratory conditions (12-h day-night cycle; lights on: 8:00-20:00 h; 22 °C) with *ad libitum* food and water unless otherwise stated. Experiments were conducted during the first half of the light phase. Rats were handled daily for a week prior to behavioral testing. All experiments were approved by the KU Leuven animal ethics committee according to European guidelines.

### Drug administration and Apparatus

Rats were intraperitoneally injected with 1 mg/kg scopolamine hydrobromide (Tocris Bioscience, category number 1414), dissolved in 0.9% NaCl solution, in a volume of 1 ml/kg. Control rats were intraperitoneally injected with saline vehicle solution. Rats were injected 30 minutes before (EXP2 and 3), or 30 minutes after (EXP3) the reinstating US presentations.

Automated operant chambers (34 cm length × 33 cm width × 33 cm height; Bilaney Consultants GmbH, Düsseldorf, Germany) were in isolation cubicles (Med Associates, Vermont, US). All chambers had metal ceilings and side walls, whereas front and back walls were clear Plexiglas. The floor was a stainless steel grid (0.5 cm in diameter), by which a constant-current foot shock could be delivered (1s, 0.4 mA). In each chamber, there was an operant lever, and adjacent to the lever was a niche. Each chamber was equipped with a water-dipper that could deliver 0.04 ml water into a cup at the bottom of the niche. Opposite the water-dipper was a nose-poke device. Rate of nose-poking was recorded with Graphic State 3.0 software (Coulbourn Instruments, Allentown, US). A speaker was mounted on the same side wall near the niche. The speaker was connected to an audio cue which produced a high frequency tone (5 kHz, 74 dB) and served as conditional stimulus (CS). The CS was presented for 30 consecutive seconds. Each chamber was illuminated by a dim house light. A camera was mounted on top of each operant chamber which recorded the rat’s activity during CS presentation at each session.

### Experimental procedure

#### Nose poke training

To motivate drinking behavior in rats, we imposed a scheduled drinking protocol, several days before the experiment. Rats were allowed a one-hour water session a day. Body weight was monitored daily to not exceed 15% initial body weight loss. Water-deprived rats were shaped daily for one hour to obtain water by using the nose-poke device. Water could be collected during all trials, but reinforcement schedules gradually increased in demand, motivating continuous response from the rats to obtain water. During days 1 and 2, rats received water every two minutes regardless of the rat’s behavior. Each nose poke was also reinforced with access to water. If the criteria of 10 nose pokes was not reached at the end of day 2, rats were hand-shaped for two additional days. During days 3 to 5, rats had to perform a nose poke to receive water access. If the criteria of 50 nose pokes in one session was reached, rats advanced to the last shaping schedule where nose pokes were reinforced on average every 20 seconds (days 6 to 9). Shaping was completed if rats sustained ≥ 120 nose pokes for three subsequent days. Rats were maintained on a VI20 schedule during the entire experiment.

#### CS acquisition

After shaping was completed, rats were subjected to a single fear acquisition session. After 15 min baseline activity, a 30 second presentation of the CS co-terminated with an electric foot shock (1s, 0.4 mA). The intertrial interval (ITI) was approximately 15 minutes and rats were exposed to four CS-shock pairings. Rats were removed from the operant chamber three minutes after the last electric foot shock.

#### Mass extinction and extinction test

Two mass extinction sessions of 20 trials each followed fear acquisition. Unreinforced CS presentations started after 10 minute baseline, with a duration of 30 s and ITI of 1 minute. Rats were removed from the operant chamber three minutes after the last CS presentation. To measure post-extinction fear memory, we presented 24 hours later four unreinforced presentations of the CS. Baseline and ITIs were on average 10 minutes.

#### Reinstatement and return-of-fear test

During the reinstatement procedure, rats received an unsignaled electric foot shock after 10 minute baseline. A second unsignaled electric foot shock was given 5 minutes afterwards. The rats remained in the operant cage for another minute before they were removed. The reinstating US presentations occurred in either the same or a different context from extinction (except in EXP3, where all rats were placed in the extinction context during reinstatement). Table 1 shows the differences between the same and different context. Twenty-four hours later, we presented four unreinforced CS presentations as a measure for return of fear in the original extinction context. Baseline and ITIs were on average 10 minutes. For an overview of the experimental design, see Table 2.

**Table 1.** Differences between the extinction context (same context) and the different context used during reinstatement.

| Original extinction context | Different context |
| --- | --- |
| Dim house light on | House light off |
| No background white noise | Background white noise |
| Chamber horizontally aligned | Chamber tilted by 5.5° |
| Same operant chamber | Different operant chamber |
| Normal odor in bedding under grid | Instanet and peppermint odor in bedding under grid |

**Table 2.** Overview of the experimental designs. Reinstatement could take place in the same or different context from extinction.

| Experiment | Day 1 | Day 2-3 | Day 4 | Day 5 | Day 6 |
| --- | --- | --- | --- | --- | --- |
|  | Acquisition | Extinction | Extinction<br>memory test | Reinstatement | Return of fear<br>test |
| EXP1 | 4 CS+ | 40 CS- | 4 CS- | Same context | 4 CS- |
|  |  |  |  | Different context |  |
| EXP2 | 4 CS+ | 40 CS- | 4 CS- | Saline-same | 4 CS- |
|  |  |  |  | Saline-diff |  |
|  |  |  |  | Scop-same |  |
|  |  |  |  | Scop-diff |  |
| EXP3 | 4 CS+ | 40 CS- | 4 CS- | Saline-pre | 4 CS- |
|  |  |  |  | Saline-post |  |
|  |  |  |  | Scop-pre |  |
|  |  |  |  | Scop-post |  |

### Data analyses

Data are expressed as means ± SEM and were analyzed with R statistical software (version 3.5.1) and JASP (version 0.9.2, JAPS team, 2019). Percentage of freezing was calculated on the number of seconds that rats remained immobile during the CS presentation and were scored by an experimenter blind to the experimental condition. For each experiment, we analyzed the acquisition data via two-way repeated measures ANOVAs with CS as within-subjects factor and Group as between-subjects factor. For the extinction and return-of-fear tests, we averaged the percentage freezing across the four trials of each test, and we analyzed each test separately via one-way (EXP1) or two-way (EXP2 and EXP3) ANOVAs. Tukey’s HSD test was used for post-hoc comparisons. In all statistical tests, p-values < .05 were considered significant.

## Results

### EXP1: Return of CS fear after reinstating US presentations is context-dependent

To assess whether reinstatement is indeed mediated by context conditioning, the reinstating US presentations took place in either the same (*N* = 12) or a different context from extinction (*N* = 12). We predicted less freezing at the subsequent return-of-fear test when the reinstating US presentations took place in the different context.

Freezing levels significantly increased during acquisition, *F*(3,66) = 42.759, *p* < .001, ◻*_p_^²^* = .660, and were similar between the groups, *F*(1,22) = 0.288, *p* = .597, ◻*_p_^²^* = .013 (Figure 1). Freezing levels were also similar between groups at the extinction test, *F*(1,22) = 1.372, *p* = .254, ◻*_p_^²^* = .059. Following the reinstating US presentations, a significant group difference emerged, *F*(1,22) = 33.615, *p* < .001, ◻*_p_^²^* = .604. These findings confirm the context-dependent reinstatement effect (Bouton, 2002).

**Figure 1.**
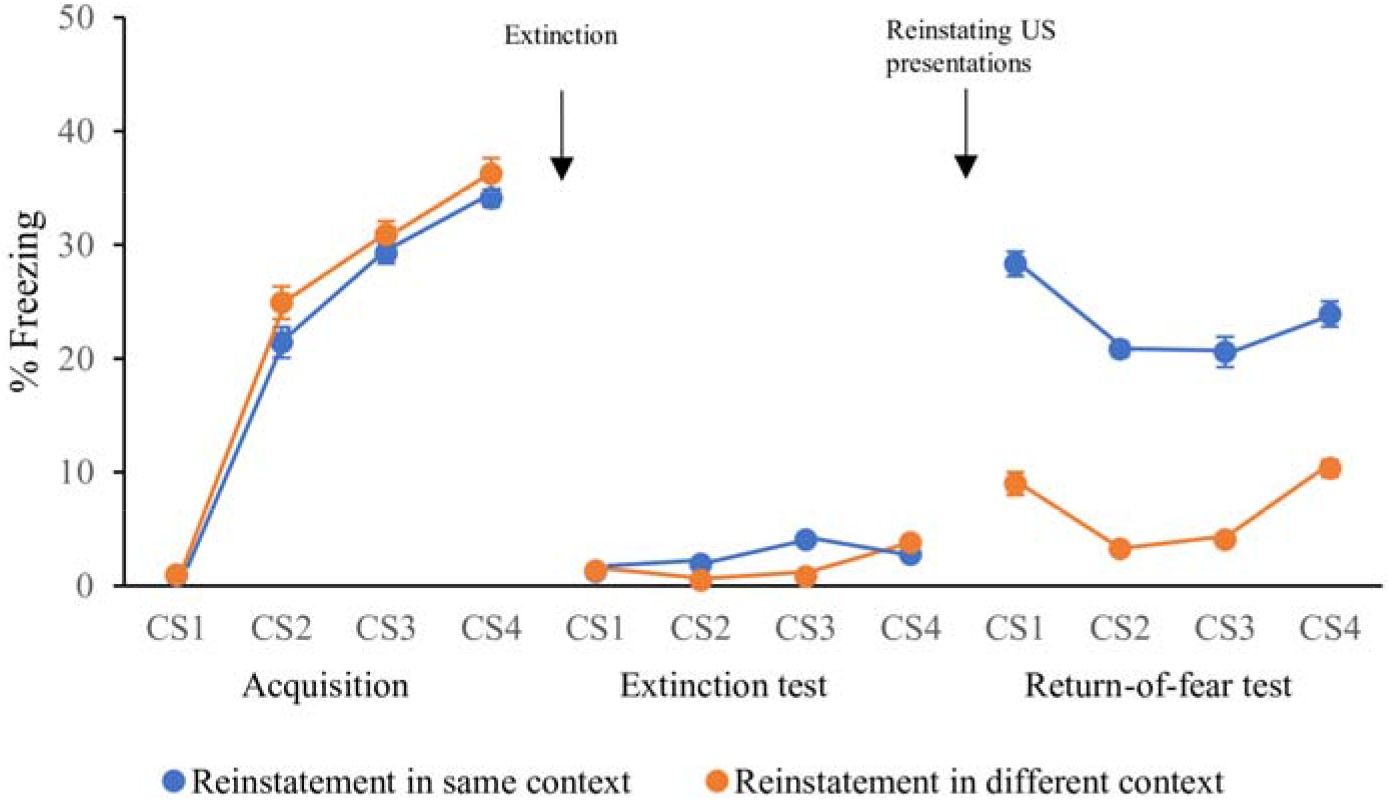
Average freezing levels during fear acquisition, the extinction test and the return-of-fear test of EXP1. During each phase of the experiment (acquisition, extinction test and return-of-fear test), the CS was presented four times. No group differences were found at fear acquisition and the extinction test. The groups differed at the return-of-fear test, with higher freezing observed when reinstatement took place in the same context. Percentage freezing on the y-axis. Error bars represent SEM. *** p < .001.

### EXP2: Systemic scopolamine before reinstating US presentations blocks context-dependent return of CS fear

Next, we evaluated whether systemic administration of scopolamine before reinstatement prevents the return of CS fear. Reinstating USs were delivered in the same or in a different context from extinction (as in EXP1), 30 minutes after scopolamine (1 mg/kg) or a vehicle solution was administered intraperitoneally, resulting in a 2 (Context: same vs different) × 2 (Treatment: saline (vehicle) vs scopolamine) design with six rats in each group (*N* = 24). We predicted less freezing at the subsequent return-of-fear test for scopolamine-treated rats (irrespective of context) and saline-treated rats tested in a different context, compared to saline-treated rats tested in the same context.

Freezing levels were similar between the groups at acquisition, as evidenced by non-significant main effects of both Context (*F*(1,20) = 0.994, *p* = .331, ◻*_p_^²^* = .047) and Treatment (*F*(1,20) = 0.846, *p* = .369, ◻*_p_^²^* = .041; Figure 2). Note that freezing during acquisition did not increase (*F*(3,60) = 1.046, *p* = .379, ◻*_p_^²^* = .050). This is because the rats were not naïve, as they had previously undergone a fear acquisition session for a different study and therefore started off with a higher level of freezing. At the extinction test, the effects of Context (*F*(1,20) = 0.396, *p* = .536, ◻*_p_^²^* = .019) and Treatment (*F*(1,20) = 0.953, *p* = .341, ◻*_p_^²^* = .046) were also non-significant, indicating that extinction memory was comparable for all groups. At the return-of-fear test, a significant interaction between Context and Treatment emerged (*F*(1,20) = 9.686, *p* = .006, ◻*_p_^²^* = .326). Post-hoc analyses revealed significantly higher freezing rates for rats that received vehicle and were presented with reinstating USs in the same context, compared to all other groups (*p*’s ≤ .003; Figure 2). No differences in freezing were observed between the other three groups (vehicle-different, scop-same and scop-different). These findings show that pre-reinstatement scopolamine administration blocks the context-dependent return of CS-evoked fear.

**Figure 2.**
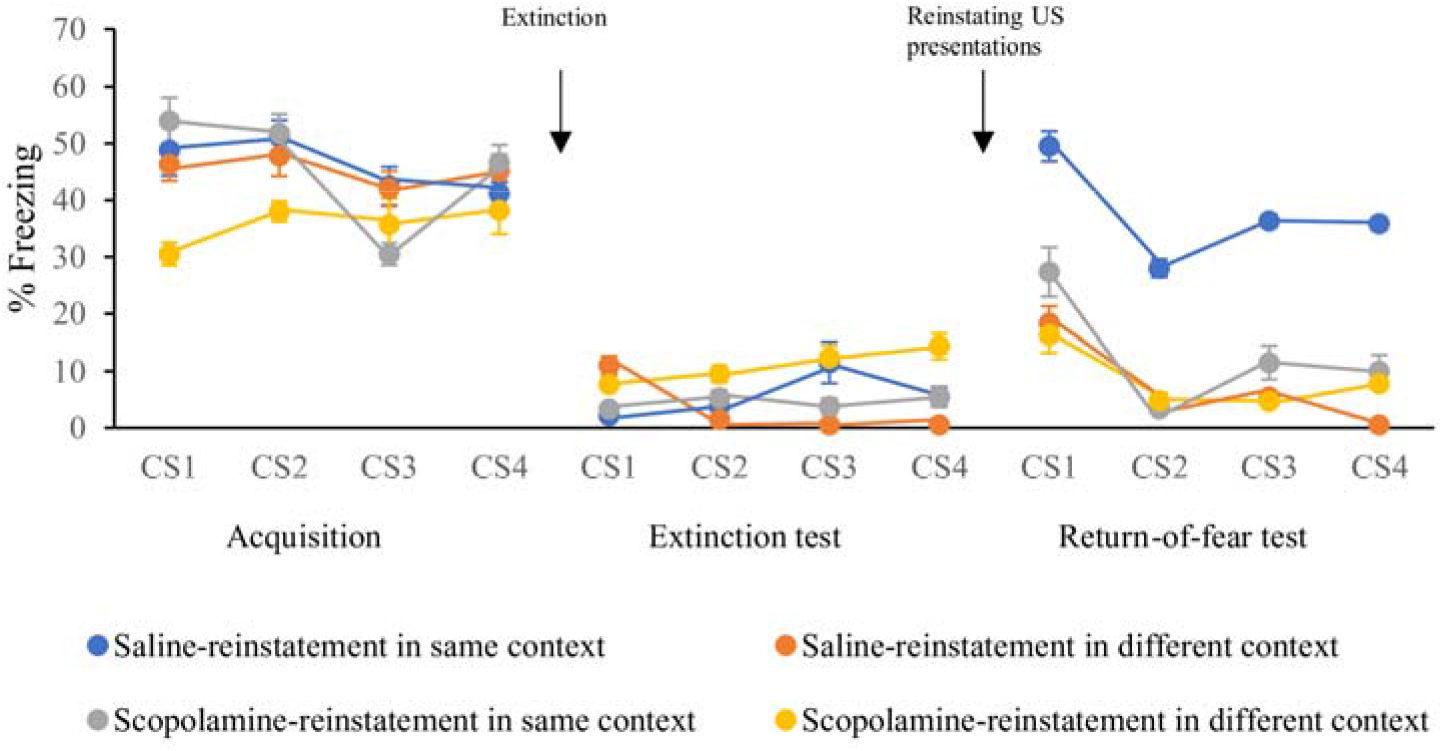
Average freezing levels during fear acquisition, the extinction test and the return-of-fear test of EXP2. During each phase of the experiment (acquisition, extinction test and return-of-fear test), the CS was presented four times. No group differences were found at fear acquisition and the extinction test. The groups differed at the return-of-fear test, with significantly higher freezing observed in the saline-same group, as compared to the other three groups. Percentage freezing on the y-axis. Error bars represent SEM. ** < .01.

### EXP3: Systemic scopolamine after reinstating US presentations blocks return of CS fear

Finally, we assessed whether post-reinstatement administration of scopolamine would also block the reinstatement effect. In this study, reinstatement took place in the same context as extinction for all rats, and scopolamine (1 mg/kg) or vehicle was administered intraperitoneally 30 minutes before or after reinstatement. This resulted in a 2 (Treatment: vehicle vs scopolamine) × 2 (Administration time: before vs after reinstatement) design with 12 rats in each group (*N* = 48). We predicted less freezing at the return-of-fear test when scopolamine was administered either before or after the reinstating US presentations. As in the previous studies, return of CS-evoked fear was measured 24 hours after reinstatement.

Freezing levels significantly increased during acquisition (*F*(3,132) = 57.737, *p* < .001, ◻*_p_^²^* = .568), and were similar between the groups, as evidenced by non-significant main effects of Treatment (*F*(1,44) = 0.224, *p* = .638, ◻*_p_^²^* = .005) and Administration time (*F*(1,44) = 0.292, *p* = .592, ◻*_p_^²^* = .007; Figure 3). At the extinction test, the effects of Treatment (*F*(1,44) = 1.991, *p* = .165, ◻*_p_^²^* = .043) and Administration time (*F*(1,44) = 0.017, *p* = .898, ◻*_p_^²^* < .001) were also non-significant. At the return-of-fear test, a significant main effect of Treatment emerged (*F*(1,44) = 130.184, *p* < .001, ◻*_p_^²^* = .747), without a Treatment × Administration time interaction (*F*(1,44) = 2.449, *p* = .125, ◻*_p_^²^* = .053). Post-hoc analyses confirmed higher freezing in the pre- and post-vehicle groups, compared to the pre- and post-scopolamine groups (*p*’s < .001). No significant differences were found between the pre- and post-vehicle group (*p* = .574) or between the pre- and post-scopolamine group (*p* = .793). This shows that the administration of scopolamine both before or after reinstatement blocks the return of CS fear.

**Figure 3.**
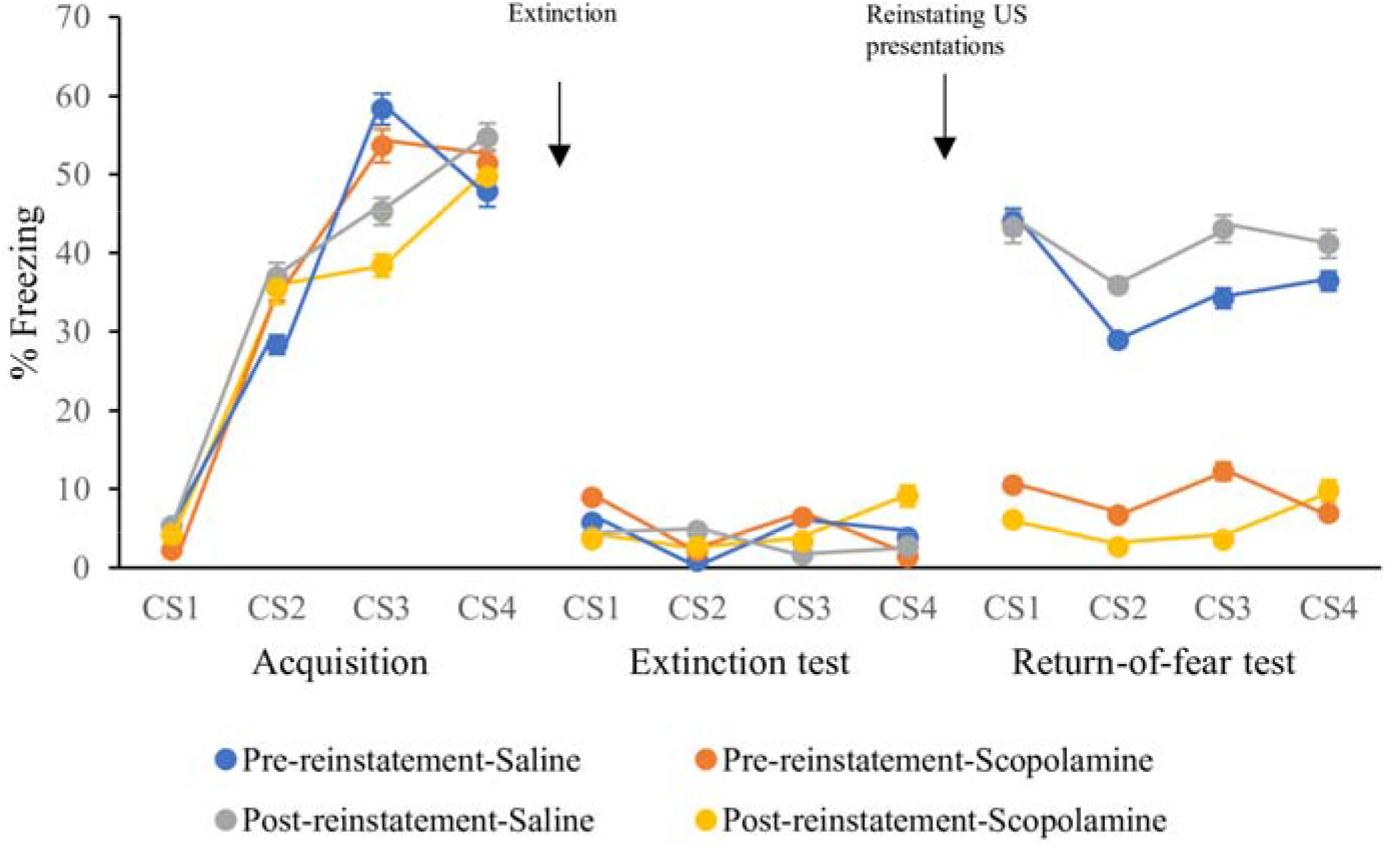
Average freezing levels during fear acquisition, the extinction test and the return-of-fear test of EXP3. During each phase of the experiment (acquisition, extinction test and return-of-fear test), the CS was presented four times. No group differences were found at fear acquisition and the extinction test. The groups differed at the return-of-fear test, with significantly higher freezing observed in the saline (both pre and post) groups, as compared to the scopolamine groups. Percentage freezing on the y-axis. Error bars represent SEM. *** p < .001.

## Discussion

The present experiments examined the effect of scopolamine on reinstatement of fear in rats. Experiment 1 confirmed the context-dependency of reinstatement. Experiments 2 and 3 showed that intraperitoneal scopolamine administration before or after reinstating US presentations blocked context-dependent return of CS fear. The present findings suggest that scopolamine impaired the context conditioning effects of reinstating US presentations that ordinarily trigger the return of CS-evoked fear.

Most pre-clinical studies have focused on optimizing fear extinction learning and consolidation to prevent later relapse effects (Walker et al., 2002; Woods & Bouton, 2006). We took a different approach by targeting reinstatement directly. Previous studies suggested that the retrieval of CS-evoked fear following reinstatement is context-dependent (Bouton & Bolles, 1979), a finding that we replicated in our first experiment. In particular, when the reinstating US presentations took place in a different context from extinction, we found no return of fear at the subsequent test. In contrast, we did observe a return of fear when reinstatement took place in the same context. These findings suggest that reinstating US presentations establish a context-US association that retrieves the CS-US association when this context is also present during the CS-alone test (Bouton, 2002). We reasoned that targeting the establishment of the context-US association during reinstatement might block the context-dependent return of CS fear.

The role of cholinergic neurotransmission in learning and memory has been well established (Klinkenberg & Blokland, 2010). Hasselmo and colleagues suggested that the cholinergic system is particularly important for the acquisition of novel information (Hasselmo & McGaughy, 2004). The system has been proposed to play a role in cortical memory encoding by enhancing afferent sensory input as well as the response of cortical circuits to such input (Hasselmo, 2006). In relation to the present study, it should be noted that the cholinergic system would therefore be particularly important in context-dependent learning. We administered scopolamine to our animals, a non-selective muscarinic antagonist, and one of the standard drugs to induce experimental amnesia (Klinkenberg & Blokland, 2010). Previous studies have repeatedly shown that pre-training systemic scopolamine blocks contextual fear conditioning in rats (Anagnostaras et al., 1995; Anagnostaras et al., 1999; Luyten et al., 2017), suggesting that inhibiting cholinergic transmission indeed affects the encoding of contextual information. Given the putative ability of scopolamine to block contextual conditioning, we evaluated its ability to prevent reinstatement as well.

In line with our *a priori* hypothesis, scopolamine administered before reinstatement prevented the return of CS-evoked fear. These results converge with previous findings that scopolamine eliminates the context-dependency of extinction (Zelikowksy et al., 2013). Similar to reinstatement, extinction is context-dependent: Fear renewal is observed when rats are presented with an extinguished stimulus in a context that is different from the one used during extinction (McNally, 2007). By administering scopolamine prior to extinction, the authors were able to block the contextual encoding that normally occurs during extinction (Zelikowsky et al., 2013), thereby preventing subsequent fear renewal. Consequently, scopolamine appears to target both fear renewal and reinstatement, by interfering with the acquisition/encoding of contextual information.

The effects of post-acquisition scopolamine administration have not been well defined (Thonnard et al., 2019). Hasselmo and colleagues proposed that cholinergic neurotransmission is differently involved in acquisition (encoding) and post-acquisition processes (Hasselmo & McGaughy, 2004). In particular, blocking cholinergic transmission would interfere with the encoding of novel information, whereas consolidation and retrieval would be more affected by stimulation of this system. Consequently, scopolamine would predominantly impair the encoding of novel information. The current study, however, yielded similar findings when scopolamine was administered before or after context conditioning during reinstatement. This suggests that scopolamine interfered predominantly with early memory consolidation. Indeed, a recent study found that scopolamine administered shortly after contextual conditioning yielded similar effects to pre-training administration, again suggesting that scopolamine impairs early memory consolidation (Luyten et al., 2017).

Our finding that scopolamine prevents the return of CS-evoked fear paves the way for new relapse prevention therapies, by thwarting context conditioning effects after aversive events. In clinical practice, impactful aversive events can reinstate previously extinguished fears and hence trigger a relapse of pathological anxiety (Vervliet et al., 2013). Scopolamine administration directly after the aversive event might prevent a subsequent return of fear, and consequently, reduce the risk of relapse. For example, a traffic victim with a history of treated panic disorder could benefit from scopolamine administration in the emergency room to prevent a revival of the old fear of panic attacks. Future studies are required to validate the use of scopolamine as a possible intervention strategy in humans. Of note, a recent clinical trial administered scopolamine to human participants in an attempt to eliminate the context-dependency of fear extinction (Craske et al., 2019). However, even though scopolamine successfully eliminates the context-dependency of fear extinction in rodents (Zelikowsky et al., 2013), the findings in humans were only limited and suggestive (Craske et al., 2019; Rothbaum & Ressler, 2019).

In summary, the current study shows that scopolamine administration at the time of reinstating US presentations effectively prevents a return of CS-evoked fear in rats. Future studies might shed more light on the effectiveness of scopolamine to prevent the establishment of a context-US association during reinstatement in humans, and the persistence of its effects over time.

## Acknowledgments

This research was supported by KU Leuven (Belgium) CREA grant CREA/13/004 and KU Leuven (Belgium) Starting Grant STG/17/035.

The authors wish to thank dr. Tom Beckers for discussions.

